# High-throughput genetics enables identification of nutrient utilization and accessory energy metabolism genes in a model methanogen

**DOI:** 10.1101/2024.03.05.583561

**Authors:** Leslie A. Day, Hans K. Carlson, Dallas R. Fonseca, Adam P. Arkin, Morgan N. Price, Adam M. Deutschbauer, Kyle C. Costa

**Affiliations:** Department of Plant and Microbial Biology, University of Minnesota, St. Paul; Environmental Genomics and Systems Biology Division, LBNL; Department of Bioengineering, University of California, Berkeley; Plant and Microbial Biology Department, University of California, Berkeley

## Abstract

Archaea are widespread in the environment and play fundamental roles in diverse ecosystems; however, characterization of their unique biology requires advanced tools. This is particularly challenging when characterizing gene function. Here, we generate randomly-barcoded transposon libraries in the model methanogenic archaeon *Methanococcus maripaludis* and use high-throughput growth methods to conduct fitness assays (RB-TnSeq) across over 100 unique growth conditions. Using our approach, we identified new genes involved in nutrient utilization and response to oxidative stress. We identified novel genes for the usage of diverse nitrogen sources in *M. maripaludis* including a putative regulator of alanine deamination and molybdate transporters important for nitrogen fixation. Furthermore, leveraging the fitness data, we provide the first evidence that *M. maripaludi*s can utilize the nitrogen sources L-glutamine, D-glucuronamide, and adenosine. Under autotrophic growth conditions, we identified a gene encoding a domain of unknown function (DUF166) that is important for fitness and hypothesize it has an accessory role in carbon dioxide assimilation. Finally, comparing fitness costs of oxygen versus sulfite stress, we identified a previously uncharacterized class of dissimilatory sulfite reductase-like proteins (Dsr-LP, group IIId) that is important during growth in the presence of sulfite. When overexpressed, Dsr-LP conferred sulfite resistance and enabled use of sulfite as the sole sulfur source. The high-throughput approach employed here allowed for generation of a large-scale dataset that can be used as a resource to further understand gene function and metabolism in the archaeal domain.

**Significance Statement:** Archaea are widespread in the environment, yet basic aspects of their biology remain underexplored. To address this, we apply randomly-barcoded transposon libraries (RB-TnSeq) to the model archaeon *Methanococcus maripaludis.* RB-TnSeq coupled with high-throughput growth assays across over 100 unique conditions identified roles for previously uncharacterized genes, including several encoding proteins with domains of unknown function (DUFs). We also expand on our understanding of carbon and nitrogen metabolism and characterize a group IIId dissimilatory sulfite reductase-like protein as a functional sulfite reductase. This dataset encompasses a wide range of additional conditions including stress, nitrogen fixation, amino acid supplementation, and autotrophy; thus, providing an extensive dataset for the archaeal community to mine for characterizing additional genes of unknown function.

## Introduction

Archaea are abundant in diverse habitats and play essential roles in ecosystem function (1) such as carbon and nitrogen cycling. Despite their ubiquity, basic aspects of archaeal physiology are still uncharacterized. To address these gaps, high-throughput methods are needed to rapidly assign putative gene function and characterize metabolic pathways.

The study of model organisms is key to overcoming the challenges associated with characterizing archaeal physiology. *Methanococcus maripaludis* is a model (2) with well-established genetic tools for understanding archaeal biology, methanogenesis, and diazotrophy. However, even in this well-studied organism, ∼35% of proteins have RefSeq annotations of either “hypothetical protein”, “membrane protein”, or belonging to a family rather than having specific functional assignments.

In the past decade the development of randomly-barcoded transposon-site sequencing (RB-TnSeq) has facilitated the rapid characterization of gene function for a variety of cellular processes in diverse microorganisms (3, 4). RB-TnSeq is a variation of TnSeq where unique DNA sequences, or barcodes, are inserted alongside transposons. Thus, RB-TnSeq can be used to conduct comparable experiments to those classically done with TnSeq, in which mutant fitness in a control condition is compared to that of an experimental condition, to identify genes with significant phenotypes. This can be done at a fraction of the effort due to the simplicity of barcode sequencing (BarSeq). Previous work has used classical transposon site sequencing to characterize the essential genes in *M. maripaludis* strain S2 (5). However, this library was challenging to use to characterize gene fitness for non-essential genes in different growth environments in high-throughput. To overcome this challenge, we developed the RB-TnSeq library which takes advantage of barcode sequencing for inexpensive detection of condition specific phenotypes.

We generated RB-TnSeq libraries for two *M. maripaludis* strains, S2 and JJ. Libraries were generated in two strains to account for a range of genotypes. The genomes of JJ and S2 are 95% identical (2-way average nucleotide identity) (6, 7), however they have major phenotypic differences. For example, strain JJ forms biofilms and can take up DNA via a natural transformation pathway in laboratory conditions while strain S2 exhibits more reliable growth in nitrogen fixing conditions. We also report methods for high-throughput cultivation and fitness profiling of these libraries to generate 536 genome-wide fitness assays representing 112 unique growth conditions, with replicates. Fitness assays were conducted across a range of autotrophic, diazotrophic, nutrient rich, and stress conditions with either H_2_ or formate as the electron donor. From this large dataset we identified novel genes of importance for carbon, nitrogen, and sulfur metabolism.

## Results

### Generation of RB-TnSeq libraries

We generated barcoded transposon libraries in both *M. maripaludis* strain S2 and strain JJ wild-type backgrounds using a barcoded variant of a mariner transposon. In this study, we focus on the analysis and discussion of the *M. maripaludis* strain S2 library as strain S2 is the model used for many aspects of archaeal biology and metabolism including determination of essential genes (5), diazotrophy (8–10), and H_2_ and formate oxidation for methanogenesis (11–13). Data from *M. maripaludis* strain JJ fitness assays was used to confirm the results from *M. maripaludis* strain S2 when feasible.

For *M. maripaludis* strain S2 we used Tn-Seq to identify 38,887 unique barcodes that mapped ≥10 times and link them to 29,150 unique insertion sites distributed across the genome (Supplemental Fig. 1). Of these barcoded insertions, 21,269 were within the central 10-90% of a gene accounting for insertions in 1,615 of the 1,741 predicted protein-coding genes. Among the genes with transposon insertions, the average number of strains per gene is 13.1, with a median of 8 strains, and a mean coverage of 284.41 reads per million mapped reads (Table 1). For strain JJ we identified 81,398 unique barcodes at 50,146 insertion locations resulting in insertions in 1,503 of the 1,796 predicted protein-coding genes (Table 1).

**Table 1.**
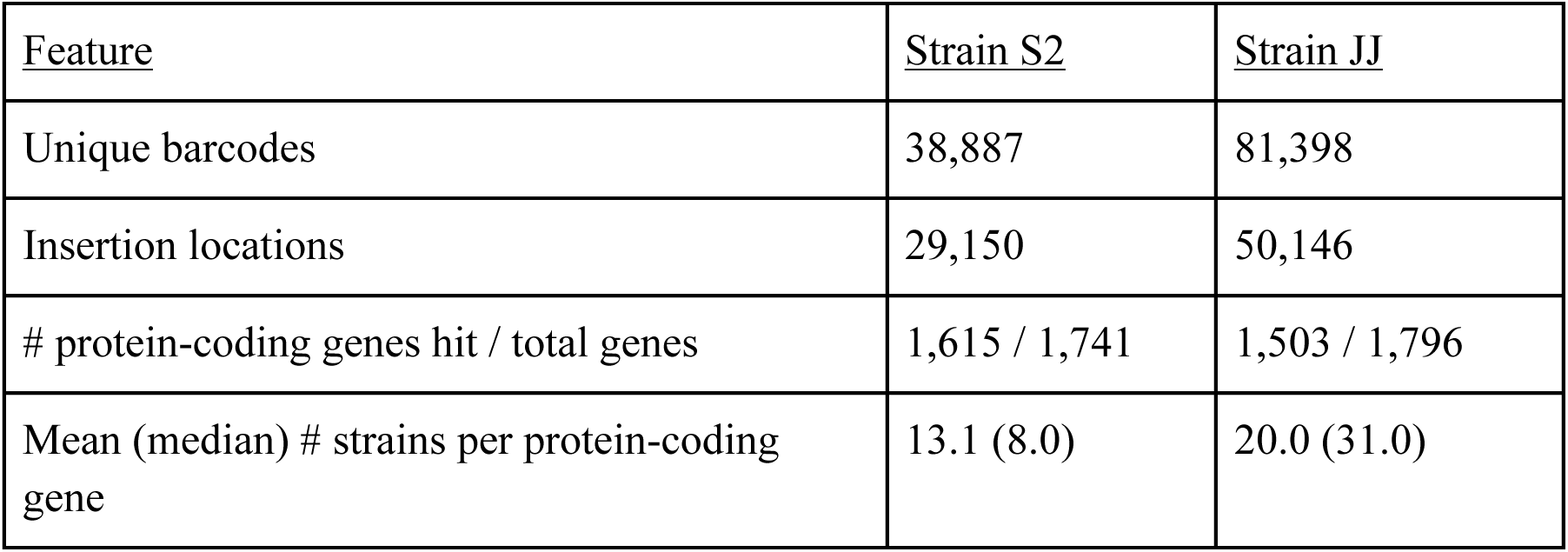
RB-TnSeq library quality statistics for *M. maripaludis* strains S2 and JJ.

| <u>Feature</u> | <u>Strain S2</u> | <u>Strain JJ</u> |
| --- | --- | --- |
| Unique barcodes | 38,887 | 81,398 |
| Insertion locations | 29,150 | 50,146 |
| # protein-coding genes hit / total genes | 1,615 / 1,741 | 1,503 / 1,796 |
| Mean (median) # strains per protein-coding gene | 13.1 (8.0) | 20.0 (31.0) |

To take advantage of the throughput and simplicity of RB-TnSeq we normalized the basal cultivation medium and optimized growth in high-throughput (Supplemental Figs. 2&3). Through incubation of 1 mL cultures in 96 well deep-well blocks in high pressure incubation vessels, we were able to conduct fitness assays in over 100 unique conditions with biological replicates. Our goal with using this approach was to conduct assays with each library in both H_2_ and formate oxidizing conditions in parallel. The only past high-throughput growth assay in *M. maripaludis* were conducted in formate oxidizing conditions (14). We calculated a normalized fitness log_2_ fold change in relative abundance of each strain. Then we calculated a weighted average of the fitness values for relevant strains for each gene and normalized gene fitness by chromosomal position (3). See materials and methods for further detail. Between the two strains and the 536 genome-wide fitness assays performed, we identified 189 genes with specific phenotypes. Specific phenotypes are defined as genes that have a strong phenotype (|fitness| > 1, |t| > 5) in a small subset of conditions tested (< 5% conditions) and lack a phenotype (|fitness| < 1) in other conditions. Henceforth we aim to highlight the phenotypes we found the most impactful and/or surprising and highlight the utility we hope this dataset will provide to the archaeal research community.

### Identification of genes important for autotrophic growth

*M. maripaludis* can fix CO_2_ to acetyl-CoA via the Wood-Ljungdahl pathway. The key enzyme complex for autotrophic growth is the carbon monoxide (CO) dehydrogenase and acetyl-CoA synthase (CODH-ACS) encoded by an operon spanning genes MMP_RS05100 to MMP_RS05065. CODH-ACS synthesizes the carbonyl group of acetyl-CoA via H_2_ and ferredoxin dependent CO_2_ reduction to CO and derives the methyl group of acetyl-CoA from an intermediate of methanogenesis (Figure 1 panel A). To determine if there were additional encoded functions essential to CO_2_ fixation, we conducted fitness assays under autotrophic growth conditions.

**Figure 1.**
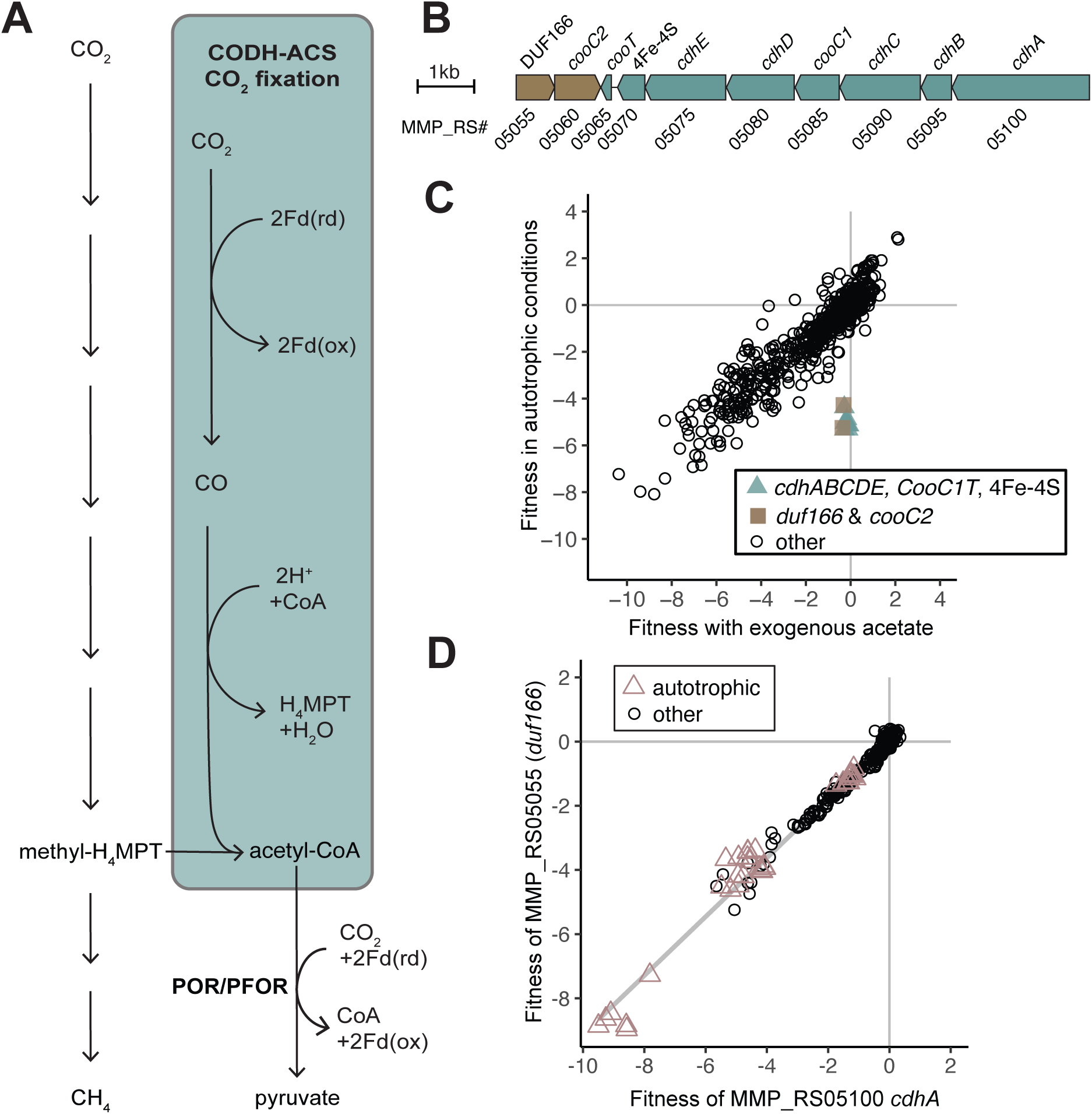
DUF166 encoded by MMP_RS05055 is important for fitness during autotrophic growth.(A) The Wood-Ljungdahl pathway for CO_2_ fixation, highlighted in blue, uses methyl-H_4_MPT from the methanogenic pathway for the methyl group of acetyl-CoA. Acetyl-CoA can be further reduced to pyruvate by pyruvate ferredoxin oxidoreductase (POR/PFOR) and reduced ferredoxin (Fd). (B) Gene neighborhood of the CODH-ACS operon. (C) Comparison of gene fitness values for growth with exogenous acetate (10 mM) compared to autotrophic growth in H_2_ oxidizing conditions. Each value is the average of 6 autotrophic (no vitamins) replicates and 3 replicates with exogenous acetate. (D) Cofitness of carbon monoxide dehydrogenase subunit alpha (*cdhA,* MMP_RS05100) and DUF166 encoded by MMP_RS05055. Experiments identified as autotrophic conditions include those both with and without vitamins.

Insertions in a cluster of 10 genes had negative fitness specifically during autotrophic growth (Figure 1 panel B&C) in H_2_ oxidizing conditions. Autotrophic growth assays were conducted in media lacking fixed carbon (casamino acids or acetate). As expected, the CODH-ACS operon accounted for 8 of the 10 genes identified. The other two genes, MMP_RS05060 and MMP_RS05055, are encoded in an adjacent operon encoded on the opposite DNA strand (Figure 1 panel B) (15). MMP_RS05060 encodes a paralog of CooC, the CO dehydrogenase nickel-insertion accessory protein thus, we will call it CooC2. (CooC1 is encoded by MMP_RS05085 in the CODH-ACS operon.) MMP_RS05055 encodes a protein with a domain of unknown function (DUF) DUF166 and is “cofit” with each gene encoded by the CODH-ACS operon and CooC2 (r = 0.94-0.99). Cofitness is defined as a linear Pearson correlation of fitness patterns (r > 0.8). The cofitness of insertions in genes encoding the CODH-ACS operon, DUF166, and CooC2 is conserved in strain JJ as well (r = 0.99-0.96).

Both MMP_RS05060 and MMP_RS05085 (in the CODH-ACS gene cluster) are annotated to encode the CooC domain predicted to be involved in maturation of the nickel center in CODH and are strongly cofit with *cdhA* (*cooC1*, r = 0.99; *cooC2*, r = 0.99). Both homologs contain the zinc CxC binding motif and conserved ADP biding residues. Additionally, the presence of both CooC domains is conserved across 340 representative genomes from different genera (16). Of the 340 genomes with both CooC homologs, 50 appear to be co-transcribed as they occur on the same strand and are within 5 kb of each other. It appears that these two CooC homologs have distinct but critical roles in CO_2_ fixation.

Insertions in DUF166 occurred on both the positive and negative strand, in both strain S2 and JJ and have negative fitness values in autotrophic conditions (Supplemental Fig. 4). However, to confirm the fitness defect was not due to a polar effect on *cooC2* we generated a gene knockout strain of *duf166* in *M. maripaludis* strain S2 (Δ*duf166*) which resulted in an inability to fix CO_2_. This defect was complemented when *duf166* was expressed in *trans* (pLW40neo-*duf166*) (Supplemental Fig. 4). This indicates the defect is due to the importance of MMP_RS05055 in autotrophic conditions and not a polar effect, suggesting its function is related to CODH-ACS carbon fixation.

The CODH-ACS complex is predicted to receive electrons from a 4Fe-4S cluster containing protein (MMP_RS05070) encoded at the end of the CODH-ACS operon (17). MMP_RS05070 is homologous to the ferredoxins PorE (39% AA ID) and PorF (37% AA ID) (Figure 2 panel A), which are encoded in an operon with the pyruvate ferredoxin oxidoreductase (POR/PFOR). In addition to being cofit with the rest of the CODH-ACS operon, MMP_RS05070 is also cofit with *porF* (MMP_RS07720), but not *porE* (MMP_RS07725) (Figure 2 panel B-D). The cofitness of MMP_RS05070 and *porF* is conserved in *M. maripaludis* strain JJ orthologs, indicating they serve a role in similar metabolic context. Additionally, the importance of *porF* in autotrophic conditions further supports previous work done using *M. maripaludis* strain JJ indicating PorF as the dominant electron conduit to POR/PFOR for pyruvate biosynthesis (18). The lack of strong fitness defects in *porE* in autotrophic conditions indicates additional experimentation is required to elucidate its role in *M. maripaludis*.

**Figure 2.**
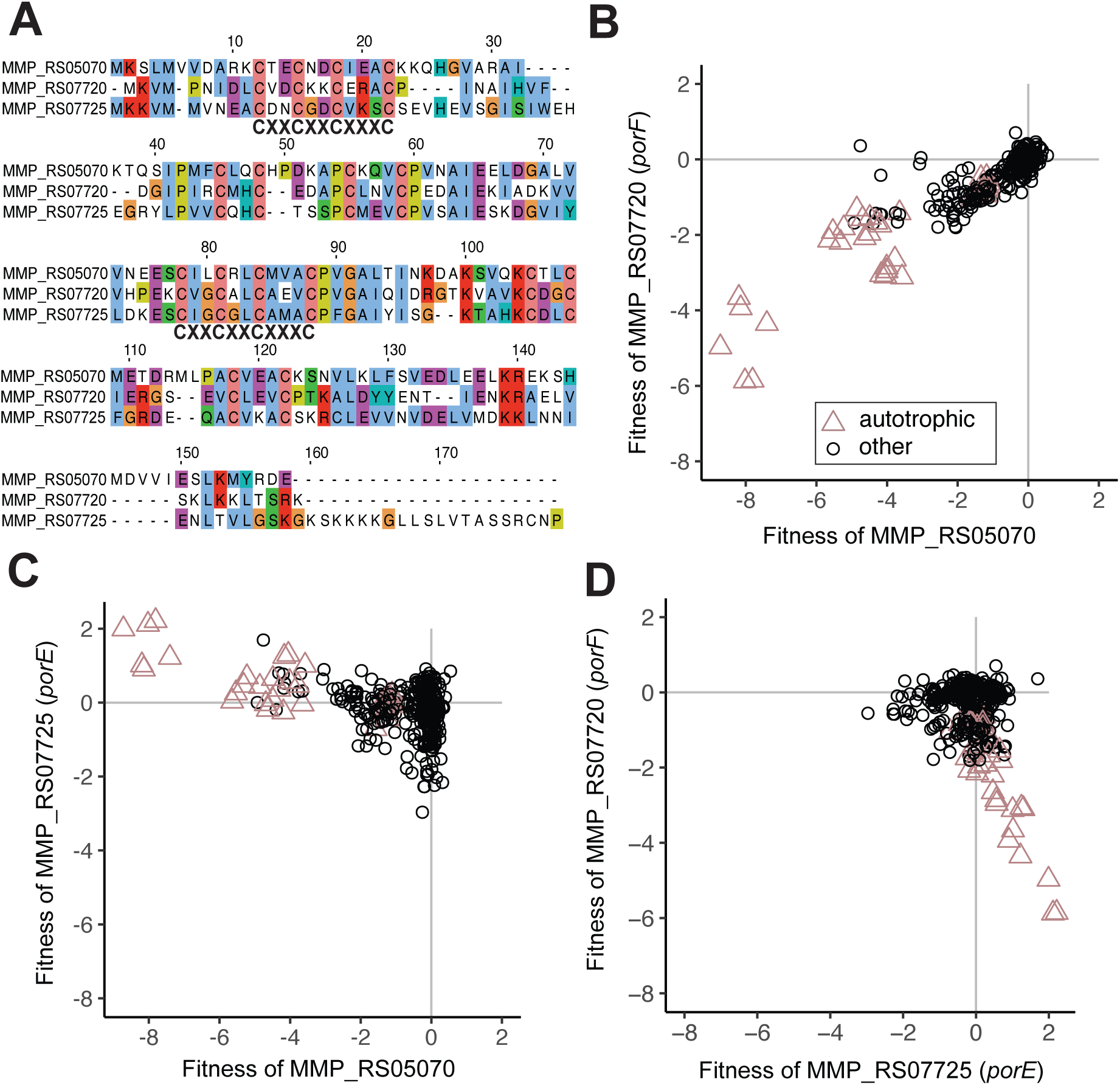
Cofitness between 4Fe-4S encoding homologs is specific to PorF and the CODH-ACS 4Fe-4S encoded by MMP_RS05070. (A) Alignment of 4Fe-FS encoding MMP_RS05070, MMP_RS07720, and MMP_RS07725 with the two conserved 4Fe-4S motifs (CXXCXXCXXXC) labeled. Amino acids are colored using the Clustal X color scheme. (B) Cofitness of MMP_RS05070 and MMP_RS07720 (PorF). (C) Negative cofitness of MMP_RS05070 and MMP_RS07725 (PorE). (C) Negative cofitness of MMP_RS07725 (PorE) and MMP_RS07720 (PorF). Experiments identified as autotrophic conditions include those both with and without vitamins.

An alternate pathway of acetyl-CoA synthesis involves assimilation of acetate from the environment via an AMP-forming acetyl-CoA synthetase (Acs). Since growth in autotrophic conditions is slower compared to growth in the presence of exogenous acetate, we would expect insertions in *acs* to have reduced fitness. However, insertions in *acs* (MMP_RS00845) have neutral fitness (|fitness| <2) across all conditions in both strain S2 and JJ. Thus, we propose the conversion of acetate to acetyl-CoA is genetically redundant between the AMP-forming acetyl-CoA synthetase, the ADP-dependent acetyl-CoA synthetase (Acd, MMP_RS01370), or possibly an acyl-CoA ligase (MMP_RS06550).

### Fitness assays in nitrogen fixing conditions

*M. maripaludis* has been studied as a model for diazotrophic growth and the nitrogenase (*nif*) operon has been well characterized (8, 9, 19, 20). Insertions in genes in the *nif* operon were detrimental in conditions where N_2_ was the sole nitrogen source provided (Figure 3 panel A). These included the molybdate dependent nitrogenase reductase (*nifH,* MMP_RS04440), nitrogenase regulators (*nifI_1,2_*, MMP_RS04445-MMP_RS04450), dinitrogenase (*nifDK,* MMP_RS04455-MMP_RS04460), and *nifEN* (MMP_RS04465-MMP_RS04470). Genes *nifHI_1,2_DKEN* are strongly cofit (r > 0.95). The last gene in the *nif* operon, *nifX* (MMP_RS04475) is non-essential in N_2_ fixing conditions as previously shown (8) (Figure 3 panel A).

**Figure 3.**
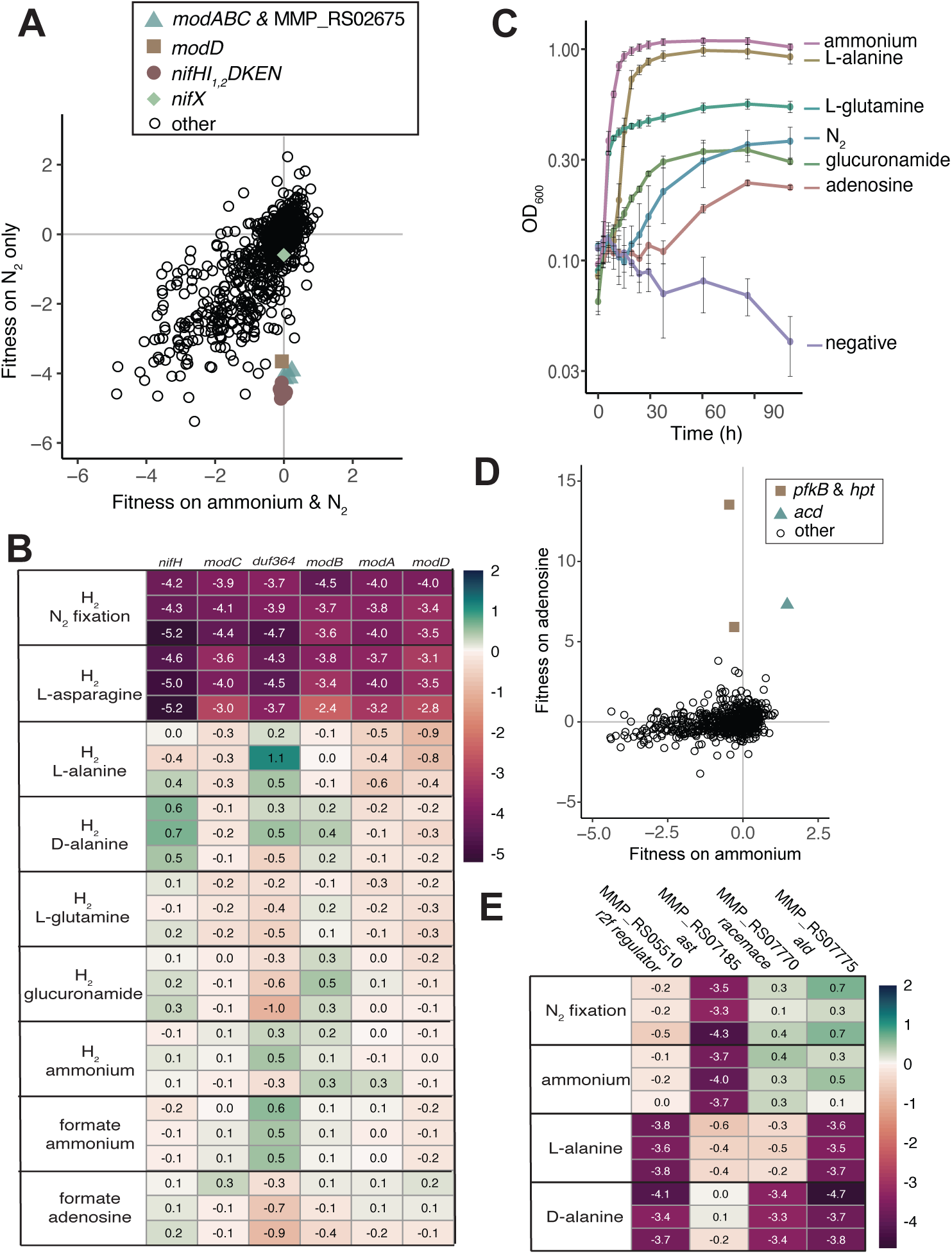
Gene fitness for diazotrophic growth and growth on various nitrogen sources. (A) Comparison of fitness values between growth with ammonium and N_2_ as nitrogen sources versus N_2_ only (both in the presence of acetate and H_2_). MMP_RS02675 encodes a DUF364. Each value is the average of 3 replicate experiments. (B) Heatmap showing overview of fitness data for select genes strongly cofit with *nifH* across conditions with varying nitrogen sources added. N_2_ is present in all conditions. (C) Growth of *M. maripaludis* with various nitrogen sources. Averages and standard error are calculated from 4 replicate cultures. N_2_ is absent in all conditions except N_2_ fixing (blue circles). (D) Fitness for growth in formate oxidizing conditions with ammonium or adenosine added as a nitrogen source. Values are averages of 3 experiments. (E) Heatmap showing fitness data for genes of interest for alanine metabolism. N_2_ is present in all conditions.

Archaeal nitrogen fixation is a molybdenum dependent process (21). *nifH* also had strong cofitness with one of the three putative molybdate transporter gene clusters (22), namely the operon comprised of *modABC* (MMP_RS02685, MMP_RS02680, MMP_RS02670) and MMP_RS02675 which encodes DUF364. Previous structural characterization and genome context analysis predicts DUF364 has a role in heavy-metal chelation (23). In organisms that encode DUF364 homologs, its presence is conserved next to molybdate uptake genes (16). We hypothesize it could have a role in chelating molybdate for biosynthesis of the molybdate cofactor (FeMoCo) required for nitrogen fixation. However, in *M. maripaludis* strain S2 insertions in MMP_RS02675 only occur on the positive strand (Supplemental Fig. 5), and we cannot rule out polar effects on downstream genes in the same predicted operon. Additionally, there is not a homolog of MMP_RS02675 in strain JJ. Thus, additional investigation is required to determine if MMP_RS02675 is required for N_2_ fixation in strain S2.

MMP_RS02730 (*modD*) is also cofit with *nifH* and is associated with another molybdate transporter encoding gene cluster *modABD* (MMP_RS02720-MMP_RS02730) previously identified as an operon (22); however recent transcriptional analysis of *M. maripaludis* (15) shows *modAB* are transcribed independent of *modD. modD* encodes a putative ATPase and its role in molybdate transport is unclear. The *modAB* encoded upstream of *modD* are non-essential and had neutral fitness across all conditions (fitness values between +/-2).The third molybdate transporter gene cluster is also comprised of two operons, *modA* MMP_RS01135 and *modBC* MMP_RS01140-MMP_RS01145 (15), is not cofit with the *nif* operon, and has weak fitness responses across conditions despite being encoded downstream of the nitrogen regulator (NrpR) operator consensus sequence GGAAN_6_TTCC (22). These results were unexpected considering past proteomic analysis performed in similar growth medium showed a significant increase in ModA encoded by MMP_RS01135 in nitrogen limiting conditions (22). Thus, under the conditions we assayed, the protein abundance of molybdate transporter subunits is not determinative of their essentiality under N_2_ fixing conditions.

### Identification of Previously Uncharacterized Nitrogen Sources

To our knowledge, N_2_, alanine, and ammonium are the only recognized nitrogen sources for *M. maripaludis* strain S2 (8, 10). We sought to identify additional nitrogen sources by screening 39 nitrogen-containing compounds in both formate and H_2_ oxidizing conditions, including 20 proteogenic amino acids as well as non-proteogenic amino acids, di-peptides, and nucleosides. To generate fitness data regardless of the substrate’s potential use as a nitrogen source, growth was performed in the presence of N_2_. In formate oxidizing conditions only cultures with D- or L-alanine, ammonium, casamino acids, L-glutamine, glucuronamide, or adenosine grew. Lack of growth in other conditions suggests that N_2_ fixation, a process that requires the H_2_-dependent reduction of ferredoxin, was poor in formate oxidizing conditions.

We used the fitness of insertions in *nifH* to determine which of the nitrogen sources were utilized, based on an approach previously implemented in *Desulfovibrio vulgaris* BarSeq analysis (24). Insertions in *nifH* had neutral fitness in H_2_ oxidizing conditions, when alanine, ammonium, L-glutamine, D-glucuronamide, or adenosine were provided as nitrogen sources, identifying L-glutamine, D-glucuronamide, and adenosine as additional nitrogen sources for *M. maripaludis* (Figure 3 panel B). The use of these substrates as nitrogen sources is further supported by growth of *M. maripaludis* in medium with L-glutamine, D-glucuronamide or adenosine as the sole nitrogen source (no N_2_ in headspace) (Figure 3 panel C).

There were no specific phenotypes when L-glutamine or D-glucuronamide were used as nitrogen sources suggesting the gene(s) responsible for cleavage of the secondary amine are essential in *M. maripaludis*, that there are redundant genes for these activities, or that hydrolysis occurs outside the cell. The three domains of the large subunit of the bacterial glutamate synthase are encoded by three essential genes in *M. maripaludis* (MMP_RS00460-MMP_RS00470). No homolog of the small subunit of the bacterial glutamate synthase has been identified (17). The lack of fitness defects for transporters in conditions where L-glutamate or D-glucuronamide is used as a nitrogen source suggests there may be redundancy in uptake mechanisms or that the transporters are encoded by essential genes.

When adenosine was provided as a nitrogen source, we identified beneficial mutations (>2 log_2_FC) for several genes predicted to be involved in purine degradation and salvage (Figure 3 panel D). Most notably, inosine kinase (*pfkB,* MMP_RS02225) had an average log_2_FC of +13.7 during growth with formate as the electron donor and adenosine and N_2_ as the sole nitrogen sources. We hypothesize that adenosine is deaminated to yield ammonium and inosine by adenosine deaminase MMP_RS07110 which is essential in *M. maripaludis* S2 (5). The inosine kinase phosphorylates inosine to IMP, costing an ATP. The other purine metabolism related gene with a strong positive selection was hypoxanthine phosphoribosyltransferase (*hpt*) MMP_RS00830. Cofitness of *hpt* and inosine kinase orthologs is conserved in *M. maripaludis* strain JJ. We hypothesize the strong positive fitness values of these genes when adenosine is used as a nitrogen source is the result of a futile cycle. Once the ammonium is cleaved from adenosine, inosine is phosphorylated to IMP which can then be phosphorylated to hypoxanthine by MMP_RS01030. Hpt then converts hypoxanthine back to inosine circularizing the futile cycle with the net effect being hydrolysis of ATP.

There was also a strong positive selection for insertions in MMP_RS01370 which is a (ADP)-forming acetyl-CoA synthetase (Acd) and hypothesized to be involved in an acetate switch (switch from acetate production to re-uptake in later growth phases) in *M. maripaludis* (25). However, *acd* being detrimental is not specific to adenosine utilizing conditions as many other conditions with acetate present and no adenosine have positive fitness values for strains with insertions in *acd*. Thus, future work is required to understand why insertions in *acd* have positive fitness values in a variety of conditions.

In all amino acid free conditions, a gene (MMP_RS07185) annotated to encode an aspartate transaminase (Ast) had negative fitness except when alanine was provided (Figure 3 panel E). Ast encodes one of five potential type I aminotransferases that could be responsible for biosynthesis of alanine (26). Alternatively, L-alanine dehydrogenase (*ald*, MMP_RS07775) is predicted to be responsible for both catabolism and biosynthesis of L-alanine (27). However, insertions in *ald* do not have strong negative fitness in amino acid free conditions except for when D- or L-alanine is provided as a nitrogen source without ammonium. Thus, we hypothesize that Ast is responsible for biosynthesis of L-alanine and Ald is likely essential for catabolism.

A Rrf2 family transcriptional regulator (MMP_RS05510) is cofit (r = 0.87) with *ald* and has negative fitness when D- or L-alanine is used as a nitrogen source (Figure 3 panel E). MMP_RS05510 does not have a strong response in other casamino free conditions, suggesting it has a regulatory role in alanine catabolism. When D-alanine is provided as a nitrogen source, insertions in the alanine racemase (MMP_RS7770) have negative fitness along with the *ald* (Figure 3 panel E), consistent with previous characterization in *M. maripaludis* (28).

### Oxygen and sulfite stress have distinct genome-wide fitness profiles

Methanogens are strict anaerobes and are therefore sensitive to a variety of oxidative stressors. Sulfite is inhibitory to methanogenesis (29) and can be produced via an abiotic reaction when oxygen is introduced to sulfide rich environments where methanogens are found (30). *In vitro,* enzymes like Methyl-coenzyme M reductase (Mcr), an essential enzyme for methanogenesis, are irreversibly damaged by sulfite (31). However, oxygen itself can also irreversibly damage metalloenzymes, so we sought to determine if there were independent consequences of oxygen and sulfite toxicity. To separate these effects, oxygen stress cultures were washed with reductant- and sulfide-free media in anoxic conditions and then agitated in air for up to one hour (to introduce oxygen, but not sulfite). The oxygen stressed cells were resuspended in anoxic media with reductant prior to the addition of sulfide, then grown in H_2_ oxidizing conditions. In contrast, the sulfite stress cultures were not exposed to air and were grown with 0.1 mM sulfite in H_2_ oxidizing conditions.

#### Increased importance of coenzyme reducing genes following oxygen exposure

There were two operons encoding genes with negative fitness in oxygen stress conditions but not control or sulfite stress conditions (Figure 4 A&B). The strongest negative fitness was for MMP_RS05965 which is annotated as a carboxymucolactone decarboxylase family protein and is encoded in a 10 gene operon (MMP_RS05955-MMP_RS06000). While other annotated carboxymucolactone decarboxylases have been characterized as disulfide reductases (MdrA) in *Methanosarcina acetivorans*, the protein encoded by *M. maripaludis* is only 30% identical to MdrA and is missing conserved cysteine residues (Supplemental Fig. 6) (32). Thus, the function of MMP_RS05965 is unclear, but is likely beneficial following oxygen exposure.

**Figure 4.**
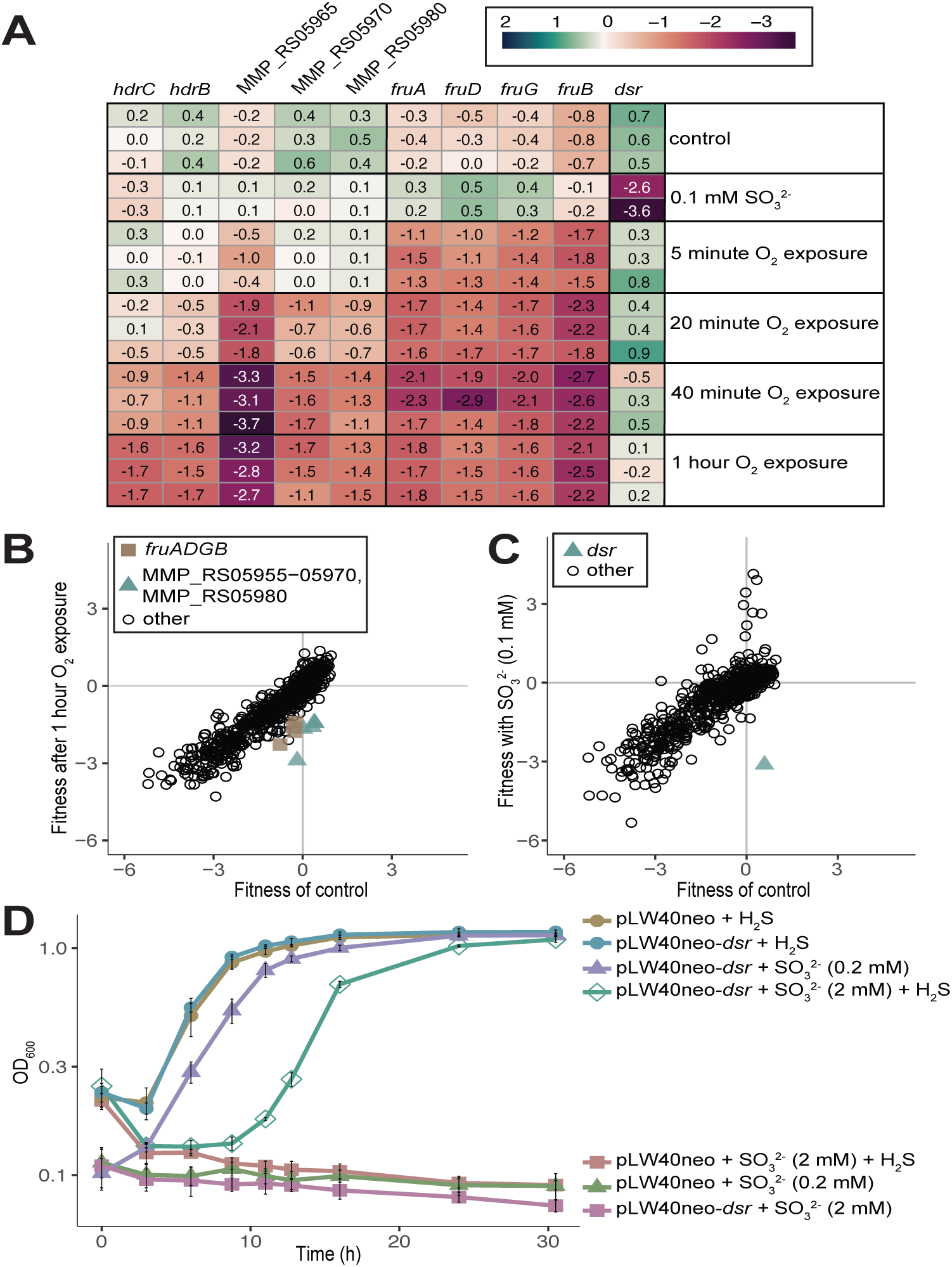
Comparison of genes with strong phenotypes in oxygen versus sulfite stress. (A) Heatmap showing fitness of select genes across control, oxygen (O_2_), and sulfite (SO_3_^2-^) stress conditions. (B) Comparison of fitness between control and oxygen exposed cultures. Values are replicates of triplicate cultures. (C) Comparison of fitness between control and sulfite containing cultures. Values are replicates of 2 cultures for sulfite and 3 replicates for control. (D) Growth of *M. maripaludis* with an empty vector (pLW40neo) or vector containing MMP_RS00450 (*dsr*) across SO_3_^2-^ stress conditions with or without sulfide (H_2_S) added as a sulfur source.

Additional genes in the operon with MMP_RS05965 have negative fitness values following oxygen exposure including those encoding heterodisulfide reductase (Hdr) subunits B and C (Figure 4 A&B). The Hdr complex is responsible for the reversible reduction of the heterodisulfide CoM-S-S-CoB to coenzymes CoM-SH and CoB-SH. A second operon with negative fitness (MMP_RS07120-MMP_RS07135) encodes the selenocysteine containing coenzyme F_420_ hydrogenase (Fru), capable of using H_2_ to reduce F_420_ which is then used by other reductive enzymes in the reduction of CO_2_ to methane (33) (Figure 4 panel A&B). Since both operons encode genes involved in redox reactions, we hypothesize that mutants have decreased fitness due to the oxidation of the coenzymes and loss of reduced pools of coenzyme F_420_H_2_ and heterodisulfide required for methanogenesis.

#### Sulfite reductase enables detoxification and use of sulfite as sole sulfur source

For testing sulfite toxicity, the transposon library was grown with 0.1 mM sulfite or 10 mM thiosulfate, both with H_2_ as the electron donor. In sulfite (Figure 4 A&C) and thiosulfate (Supplemental Fig. 7) stress conditions, strains with insertions in MMP_RS00450 had negative fitness compared to controls (Figure 4 A&C). MMP_RS00450 encodes a gene predicted to be a group IIId dissimilatory sulfite reductase-like protein (Dsr) (34).

Sulfide is the only known sulfur source for *M. maripaludis* (35). However, some close relatives with a greater sulfite tolerance are able to use sulfite through reduction to sulfide by an F_420_- dependent sulfite reductase (Fsr) (31). Expression of the Fsr from *Methanocaldococcus jannashii* in *M. maripaludis* confers the ability to reduce sulfite to sulfide for both detoxification and use as a sulfur source (36). This sulfite reductase is a chimera of a N terminal homolog of F_420_H_2_ oxidase (Fsr-N) and a C terminal homolog of Dsr (Fsr-C). However, *M. maripaludis* does not contain this chimeric form of a sulfite reductase.

The *M. maripaludis* Dsr (MMP_RS00450) contains a siroheme and two 4Fe-4S clusters but no F_420_H_2_ oxidase (Fsr-N) fusion (34, 37). However, decreased mutant fitness for the native *dsr* in *M. maripaludis* fitness assays with sulfite and thiosulfate suggests it still has a role in sulfite detoxification. We recombinantly expressed MMP_RS00450 under the control of the high expression promoter (phmv; histone promoter from *Methanococcus voltae*) (9) in *M. maripaludis* to investigate if increased expression of the native *dsr* was sufficient to confer increased sulfite resistance and use as the sole sulfur source (Figure 4D). *M. maripaludis* with recombinantly expressed Dsr was able to use 0.2 mM sulfite as the sole sulfur source. The overexpression strain was also able to detoxify 2 mM sulfite but not as the sole sulfur source (Figure 4D).

Sulfite and thiosulfate concentrations are typically in the low micromolar range or below in marine sediments (38). The Dsr-LP in *M. maripaludis* is functional as a sulfite reductase despite lacking the F_420_H_2_-oxidase domain, albeit at lower concentrations than that previously demonstrated (2 mM) with recombinant expression of *M. jannashii* Fsr (36). Since the F_420_H_2_-oxidase domain is lacking, we predict the electron donor for sulfite reduction is either essential or redundant in *M. maripaludis*; however further characterization is required to identify the electron donor for this reaction. Taken together, we hypothesize that Dsr-LP activity, while less robust than Fsr, may still be beneficial to *Methanococcus* spp. for the detoxification of sulfite under physiologically relevant conditions in anoxic marine environments.

## Conclusion

By combining RB-TnSeq with high-throughput cultivation, we successfully assessed fitness of non-essential genes in the model archaeon *M. maripaludis* in over 100 unique growth conditions. This led to identifying several genes encoding proteins of unknown function as important in diverse cellular processes. Further, we were able to characterize three new nitrogen sources, a sulfur source, and identify different genes important for the response to oxygen or sulfite stress. Finally, the determination of a functional *dsr* in *M. maripaludis* provides novel insight to sulfur metabolism for *M. maripaludis* and is the first *in vivo* analysis of the function of a Dsr-LP group IIId protein as a sulfite reductase.

RB-TnSeq is particularly valuable when *in silico* gene functional assignments are incorrect or missing. The data presented here and available through the fitness browser (http://fit.genomics.lbl.gov/) are a valuable resource for further hypothesis generation with regards to gene function. We have highlighted a handful of findings from this dataset but there are 189 genes with specific phenotypes and additional cofitness and conserved cofitness patterns to be explored. As the first application to a member of the Archaea, this RB-TnSeq dataset will empower future efforts to characterize novel aspects of archaeal physiology.

## Materials and Methods

### Growth Conditions

All *M. maripaludis* growth plates and deep-well blocks (Costar) were inoculated and harvested in an anaerobic chamber (Coy Laboratory Products, 2-3% H_2_, 10% CO_2_, N_2_ balanced atmosphere). Plates were incubated at 37°C with 10 to 20 ml of 25% Na_2_S on a paper towel in a falcon tube or plate as a sulfur source and under a pressurized atmosphere of 140 kPa H_2_:CO_2_ (80:20). For 96 well deep-well block experiments, anoxic H_2_O was added to the Na_2_S soaked paper towel each time the incubation vessels were opened to increase the humidity and reduce evaporation.

The 1000x trace mineral solution used for library generation and growth was prepared by making 21 g/L Na_3_Citrate·2H_2_0 solution and adjusting the pH to 6.5, then in order 5 g/L MnSO_4_·2H_2_O, 1 g/L CoCl_2_·6H_2_O, 1 g/L ZnSO_4_·7H_2_O, 0.1 g/L CuSO_4_·5H_2_O, 0.1 g/L AlK(SO_4_)_2_, 0.1 g/L H_3_BO_3_, 1 g/L Na_2_MoO_4_·2H_2_O, 0.25 g/L NiCl_2_·6H_2_O, 2 g/L Na_2_SeO_3_, 0.1 g/L VCl_3_, 0.033 g/L Na_2_WO_4_·2H_2_O was added. Aliquots (50 mL) of the trace mineral stock solution were bubbled under N_2_ for 20 minutes and stored in the dark in serum bottles sealed with butyl stoppers at 4°C.

For fitness assays and nitrogen fixation growth curves 200 mM MOPS buffer (pH 7.2) was added to increase the buffering capacity and reduce the differences between formate and H_2_ growth medium. Trace minerals with SO_4_^2-^ anions were swapped for Cl^-^ anion alternatives and all Mg was provided as MgCl_2_ to eliminate SO_4_^2-^ for future SO_4_^2-^ free experiments. Fitness assays to determine if there were genes with specific phenotypes in the medium with MOPS added or SO_4_^2-^ removed compared to the medium used to generate the library were performed (Supplemental Fig. 2).

Balch tubes were pressurized with 280 kPa H_2_:CO_2_ (80:20) or 207 kPa N_2_:CO_2_ (80:20) with either H_2_ or formate oxidizing medium. With the exception of N_2_ fixation growth tubes which were pressurized to 103 kPa N_2_:CO_2_ then brought up to 280 kPa with H_2_:CO_2_ and repressurized with H_2_:CO_2_ daily.

### Plasmid construction

All strains, constructs, and primers used can be found in Supplemental Tables 1 and 2. For construction of the barcoded plasmid for generation of the RB-TnSeq libraries plasmid, pJJ605 (39) was digested with NotI at 37°C for 20 minutes then BstB1 was added and incubated at 65°C for an additional 20 minutes. The insertion fragment included DNA between the NotI and BstB1 restriction sites, the U1 U2 priming sites, and the 20 bp DNA barcode (3). Oligos 109-pLD26_insert1 and 110-pLD26_insert2 were used as template DNA for amplification using 107-pLD26_F and 108-pLD26_R for 10 replicate reactions to minimize bottlenecks for barcode diversity (Supplemental Table 2). PCR reaction conditions were 98°C for 1 min followed by 30 cycles of 98°C for 20s, 56°C for 20s, and 72°C for 30s, followed by a final extension at 72°C for 5 min. The 10 reactions were then pooled and aliquoted into 10 equal volumes for purification. The pLW40neo construct is a stable plasmid used for expression of genes under the control of the *Methanococcus voltae* histone promoter in *M. maripaludis* (9). The plasmid used for complementation of the ΔMMP_RS05055 (*duf166*) strain was generated by amplifying MMP_RS05055 using oligos 156-DUF166F and 157-DUF166R and Gibson assembling (40) the product into pLW40neo which had been digested with NsiI and AscI. Similarly, the gene encoding the dissimilatory sulfite reductase-like protein (*dsr*, MMP_RS00450); was PCR amplified using oligos 142-DsrF and 143-DsrR and Gibson assembled (40) into pLW40neo which had been digested with NsiI and AscI.

Digested pJJ605 and all PCR products were purified using Invitrogen PCR cleanup kit (REF K310001) per manufacturer’s protocol. Plasmid and insert fragment(s) were ligated using Gibson assembly with NEBuilder HiFi DNA Assembly Master Mix (CAT E2621) per manufacturer’s guidelines and electroporated into electrocompetent DH5alpha *E. coli*. For the barcoded pJJ605 construct (pLD026), electroporation reactions were pooled and recovered overnight, shaking at 37°C in 500 mL lysogeny broth with kanamycin (50 ug/mL) and ampicillin (50 ug/mL). *E. coli* containing pLW40neo-*dsr* was selected for on LB agar plates with ampicillin (50 ug/mL). All plasmids were purified using the Invitrogen PureLink HiPure Plasmid Filter Maxiprep Kit (REF K210016) per manufacturer’s guidelines. All insertion fragments were amplified using Phusion HF polymerase.

### Transposon library generation and RB-TnSeq

Transposon library mutagenesis was performed following the optimized protocol described by Fonseca *et. al.* (7, 39, 41) with slight differences in the starting culture preparation. Briefly, 16 overnight cultures in stationary phase (OD_600_ ∼1.0) were supplemented with 1 mL fresh McCas and repressurized to 276 kPa H_2_:CO_2_ and incubated shaking at 37°C for 2 hours. While cultures were growing, the transposon reaction was set up in the reaction buffer with pLD026 and *HimarI* transposase then incubated at 28°C for 2 hours. Transposon reactions were transferred into wild-type *M. maripaludis* using the PEG transformation method for 16 replicate cultures (42) followed by a 4 hour outgrowth. The replicate transformations were pooled and transformants were selected on McCas agar medium with puromycin and grown at 37°C for 4 days in an anaerobic incubation vessel pressurized to 138 kPa H_2_:CO_2_. Plates were flooded with 2 mL 20% glycerol McCas solution (5) and scraped into a sterile collection vessel then single use 1 mL aliquots were distributed into serum bottles and flash frozen with liquid nitrogen and stored at -80°C.

Genomic DNA was extracted using the Qiagen Blood and Tissue Kit (CAT 69504) per manufacturer’s guidelines and sequenced at the University of Minnesota Genomics Center. The genomic locations and associated barcodes were determined using a previously described Tn-Seq protocol (3). We also mapped transposon insertions and linked them to their associated DNA barcodes using a two-step PCR approach with a splinkerette adapter design (AMD, in preparation), similar to described by Rubin *et al.* (43). Fitness experiments were only included in subsequent analysis if they passed previously established quality metrics (3).

### RB-TnSeq Fitness Assays

For reviving the library, a 1 mL aliquot was thawed in the anoxic chamber and inoculated into 50 mL McCas with sulfide in a serum bottle and pressurized to 138 kPa H_2_:CO_2_. Cultures were incubated overnight then centrifuged and resuspended in anoxic basal medium (unless otherwise indicated) twice at 4000 rcf and resuspended in Basal Mc. The optical density (OD_600_) was determined on a Spectronic20, blanked with Basal Mc. After washing, for each experiment at least 3 time zero samples, 1 mL each, were collected by centrifuging at 15,000 rpm in microcentrifuge tubes and stored at -80°C. The remaining culture was then diluted to an estimated OD_600_ of 0.04 and added to 2x media for a starting density of 0.02 Abs. Cells were washed and resuspended in media type according to the assay being performed. The full list of experiments performed for both mutant libraries along with detailed metadata are available on Figshare (https://doi.org/10.6084/m9.figshare.24195057.v1).

Media and anoxic water used for deep-well blocks was prepared by bubbling with N_2_:CO_2_ or N_2_ for 30 minutes before transferring to the anoxic chamber for 1-2 days before addition of reductant (DTT), sulfide (1 mM) and filter sterilization. When additional components were added to the media as indicated in Compound_1 and Compound_2 columns of the fitness assay description, they were filter sterilized and added in a 2x concentration to 96 well deep-well blocks and covered with a AeraSeal^TM^ gas permeable film (Millipore Sigma) then transferred into the anoxic chamber to degas for 2 days before inoculating with cells in 2x media containing reductant and transferring plates to the incubation vessel. Growth of cultures was periodically checked by transferring 150 uL of culture to a Greiner 96 well microplate using an Avidien microPro 300 Benchtop pipettor. The OD_600_ was read inside the anoxic chamber using a Tecan Infinite^®^ 200 plate reader, then cultures were returned to the deep-well blocks.

### Sulfite and Oxidative Stress Assay

The sulfite stress assay cells were washed with McCas without DTT or sulfide. Culture was divided and resuspended in a 2x buffer that contained DTT and inoculated into deep-well blocks. Culture reserved for the oxygen stress assay was resuspended in media without DTT, removed from the anaerobic chamber, and incubated shaking in a flask at 37°C with foil loosely over the top. 1 mL of culture was collected after 5, 20, 40 and 60 minutes of oxygen exposure. The culture was centrifuged for 1 min at 8000 x g and the supernatant discarded and then cycled into the anoxic chamber where it degassed for 5 minutes before 2x media containing reductant (DTT) was used to resuspend the pellet. Cells remained in the 2x containing McCas for at least 10 minutes before being transferred to deep-well blocks containing sulfide.

### Syntrophic fitness assays

Fitness assays conducted with *Syntrophotalea carbinolica* were performed as previously described except with the S2 barcoded transposon library (44).

### Vitamin fitness assays

For vitamin dropout conditions a filter sterilized vitamin solutions were made lacking one of the vitamins for each vitamin present in the stock solutions (e.g. biotin dropout condition). A solution of all 20 amino acids (1 mM) each was added in place of casamino acids to ensure excess of all amino acids (formate McAA). However, there were few outliers from controls with complete vitamin solutions likely due to residual intracellular pools being sufficient to complement defects thus future studies will require serial transfers.

### Data Analysis

Essential genes have been previously characterized in *M. maripaludis* strain S2 using a derivative of the *Tn*5 transposon library (5). Similar to what was observed in the *Tn*5 transposon library, we discovered that insertions in many of these essential genes were present in the RB-TnSeq library, likely due to the polyploidy of *M. maripaludis* (45). The fitness data for essential genes complicates the normalization by chromosomal position, which is based on the assumption that most genes have little effect on fitness (3). Additionally, essential genes have a higher average GC content compared to non-essential genes (35% vs 33% respectively) which impedes determination of the quality metric (gccor) which accounts for PCR-based spurious GC content correlations (3). Therefore, we removed fitness data for insertions in 300 of the previously determined essential genes which had consistently negative fitness across conditions (without normalizing by location on the chromosome) based on unnormalized values, resulting in the exclusion of 300 genes (Supplemental Dataset and Figshare https://doi.org/10.6084/m9.figshare.24195057.v1).

Gene fitness was calculated as described previously (3). Briefly, insertions in a gene were considered if they occurred within the middle 10-90% of the gene. Strain fitness values are calculated from the normalized log_2_ ratios of the abundance of the barcode following selection versus in the starting population before selection (Time0). Gene fitness is then calculated as the weighted average of the fitness of the individual mutant strains then normalized based on chromosome position to correct for variation in copy number along the chromosome using the running median along the chromosome (excluding essential genes values for strain S2). Finally, gene fitness is normalized so that the mode is zero. Specific phenotypes are identified for genes in an experiment where |fitness| > 1 and |t| > 5 with |fitness| <1 in at least 95% of total experiments, and that the fitness value of the outlier experiment(s) were more pronounced than most other experiments conducted (|fitness|>95th percentile(|fitness|) + 0.5) (4). Cofitness was calculated as the Pearson correlation coefficient (*r*) across all fitness assays for each pair of genes. Gene pairs were considered cofit if they had a *r* > 0.8 and conserved cofitness for gene pairs included genes where cofitness is greater than 0.6 for the two genes and cofitness > 0.6 for two orthologous genes.

### Data Availability

Data for successful experiments is available via the Fitness Browser (http://fit.genomics.lbl.gov) which includes details of the experimental conditions, quality metrics for each experiment, per-strain fitness values, gene fitness scores, cofitness and conserved cofitness, and *t* values. All data from the fitness browser and tab-delimited files for *M. maripaludis* strains S2 and JJ are also archived on Figshare at https://doi.org/10.6084/m9.figshare.24195057.v1. A list of essential genes excluded from the fitness browser a figshare can be found in the Supplemental Dataset. All data from the Fitness Browser as of February 2024 is archived on Figshare at https://doi.org/10.6084/m9.figshare.25236931.v1.

## Acknowledgements

We thank Jon Badalamenti and Catalina Vega Hurtado for preparation of the DNA libraries for TnSeq. This work was supported by the U.S. Department of Energy, Office of Science, Basic Energy Sciences under grant number DE-SC0019148. This material was supported by the U.S. Department of Energy Graduate Student Research (SCGSR) program fellowship to LAD. The SCGSR program is administered by the Oak Ridge Institute for Science and Education for the U. S. Department of Energy under contract number DE-SC0014664. Work by HKC, APA, MNP, and AMD was supported by ENIGMA - Ecosystems and Networks Integrated with Genes and Molecular Assemblies (http://enigma.lbl.gov), a Science Focus Area Program at Lawrence Berkeley National Laboratory is based upon work supported by the U.S. Department of Energy, Office of Science, Office of Biological & Environmental Research under contract number DE-AC02-05CH11231.

## Notes

### Competing Interest Statement

The authors have declared no competing interest.

## REFERENCES

1. P. Offre, A. Spang, C. Schleper, Archaea in biogeochemical cycles. Annu. Rev. Microbiol. 67(0), 437–457 (2013).

2. K. C. Costa, W. B. Whitman, Model organisms to study methanogenesis, a uniquely archaeal metabolism. J. Bacteriol. 205(0), e0011523 (2023).

3. K. M. Wetmore, et al., Rapid quantification of mutant fitness in diverse bacteria by sequencing randomly bar-coded transposons. mBio 6(0), e00306–15 (2015).

4. M. N. Price, et al., Mutant phenotypes for thousands of bacterial genes of unknown function. Nature 557(7706), 503–509 (2018).

5. F. Sarmiento, J. Mrazek, W. B. Whitman, Genome-scale analysis of gene function in the hydrogenotrophic methanogenic archaeon *Methanococcus maripaludis*. Proc. Natl. Acad. Sci. U.S.A. 110(12), 4726–4731 (2013).

6. L. M. Rodriguez-R, K. T. Konstantinidis, The enveomics collection: a toolbox for specialized analyses of microbial genomes and metagenomes. PeerJ (PeerJ Inc., 2016). 10.7287/peerj.preprints.1900v1 (January 27, 2024)

7. D. R. Fonseca, et al., Random transposon mutagenesis identifies genes essential for transformation in *Methanococcus maripaludis*. Mol. Genet. Genomics 298(3), 537–548 (2023).

8. P. S. Kessler, J. A. Leigh, Genetics of nitrogen regulation in *Methanococcus maripaludis*. Genetics 152(3), 1343–1351 (1999).

9. J. A. Dodsworth, J. A. Leigh, Regulation of nitrogenase by 2-oxoglutarate-reversible, direct binding of a PII-like nitrogen sensor protein to dinitrogenase. Proc. Natl. Acad. Sci. U.S.A. 103(26), 9779–9784 (2006).

10. T. J. Lie, J. A. Leigh, Regulatory response of *Methanococcus maripaludis* to alanine, an intermediate nitrogen source. J. Bacteriol. 184(19), 5301–5306 (2002).

11. K. C. Costa, et al., Protein complexing in a methanogen suggests electron bifurcation and electron delivery from formate to heterodisulfide reductase. Proc. Natl. Acad. Sci. U.S.A. 107(25), 11050–11055 (2010).

12. K. C. Costa, T. J. Lie, M. A. Jacobs, J. A. Leigh, H_2_-independent growth of the hydrogenotrophic methanogen *Methanococcus maripaludis*. mBio 4(2), e00062–13 (2013).

13. S. T. Lohner, J. S. Deutzmann, B. E. Logan, J. Leigh, A. M. Spormann, Hydrogenase-independent uptake and metabolism of electrons by the archaeon *Methanococcus maripaludis*. ISME J 8(8), 1673–1681 (2014).

14. M. R. Weimar, et al., Development of multiwell-plate methods using pure cultures of methanogens to identify new inhibitors for suppressing ruminant methane emissions. Appl. Environ. Microbiol. 83(22), e00396–17 (2017).

15. W. Zhang, et al., Internal transcription termination widely regulates differential expression of operon-organized genes including ribosomal protein and RNA polymerase genes in an archaeon. Nucleic Acids Res. 51(14), 7851–7867 (2023).

16. M. N. Price, A. P. Arkin, A fast comparative genome browser for diverse bacteria and archaea. (2023).

17. E. L. Hendrickson, et al., Complete genome sequence of the genetically tractable hydrogenotrophic methanogen *Methanococcus maripaludis*. J. Bacteriol. 186, 6956–6969 (2004).

18. W. Lin, W. B. Whitman, The importance of PorE and PorF in the anabolic pyruvate oxidoreductase of *Methanococcus maripaludis*. Arch Microbiol 181, 68–73 (2004).

19. P. S. Kessler, C. Blank, J. A. Leigh, The *nif* gene operon of the methanogenic archaeon *Methanococcus maripaludis*. J. Bacteriol. 180, 1504–1511 (1998).

20. T. J. Lie, J. A. Leigh, A novel repressor of *nif* and *glnA* expression in the methanogenic archaeon *Methanococcus maripaludis*. Mol Microbiol 47, 235–246 (2003).

21. P. S. Kessler, J. McLarnan, J. A. Leigh, Nitrogenase phylogeny and the molybdenum dependence of nitrogen fixation in *Methanococcus maripaludis*. J. Bacteriol. 179, 541– 543 (1997).

22. Q. Xia, et al., Quantitative proteomics of nutrient limitation in the hydrogenotrophic methanogen *Methanococcus maripaludis*. BMC Microbiol. 9, 149 (2009).

23. M. D. Miller, et al., Structure of the first representative of Pfam family PF04016 (DUF364) reveals enolase and Rossmann-like folds that combine to form a unique active site with a possible role in heavy-metal chelation. Acta Crystallogr Sect F Struct Biol Cryst Commun 66, 1167–1173 (2010).

24. V. V. Trotter, et al., Large-scale genetic characterization of the model sulfate-reducing bacterium, *Desulfovibrio vulgaris* Hildenborough. Front Microbiol. 14, 1095191 (2023).

25. C. H. Vo, N. Goyal, I. A. Karimi, M. Kraft, First observation of an acetate switch in a methanogenic autotroph (*Methanococcus maripaludis* S2). Microbiol Insights. 13, 1–7 (2020).

26. R. Saum, A. Mingote, H. Santos, V. Müller, A novel limb in the osmoregulatory network of *Methanosarcina mazei* Gö1: Nε-acetyl-β-lysine can be substituted by glutamate and alanine. Environ. Microbiol. 11, 1056–1065 (2009).

27. N. Goyal, Z. Zhou, I. A. Karimi, Metabolic processes of *Methanococcus maripaludis* and potential applications. Microb Cell Factories. 15, 107 (2016).

28. B. C. Moore, J. A. Leigh, Markerless mutagenesis in *Methanococcus maripaludis* demonstrates roles for alanine dehydrogenase, alanine racemase, and alanine permease. J Bacteriol 187, 972–979 (2005).

29. W. L. Balderston, W. J. Payne, Inhibition of methanogenesis in salt marsh sediments and whole-cell suspensions of methanogenic bacteria by nitrogen oxides. Applied and Environ Microbiol. 32, 264–269 (1976).

30. J. D. Cline, F. A. Richards, Oxygenation of hydrogen sulfide in seawater at constant salinity, temperature, and pH. Environ Sci Technol. 3(9), 838–843(1969).

31. E. F. Johnson, B. Mukhopadhyay, A new type of sulfite reductase, a novel coenzyme F_420_-dependent enzyme, from the methanarchaeon *Methanocaldococcus jannaschii*. JBC 280, 38776–38786 (2005).

32. D. J. Lessner, J. G. Ferry, The archaeon *Methanosarcina acetivorans* contains a protein disulfide reductase with an iron-sulfur cluster. J Bacteriol 189, 7475–7484 (2007).

33. E. L. Hendrickson, J. A. Leigh, Roles of coenzyme F420-reducing hydrogenases and hydrogen- and F_420_-dependent methylenetetrahydromethanopterin dehydrogenases in reduction of F_420_ and production of hydrogen during methanogenesis. J. Bacteriol. 190, 4818–4821 (2008).

34. D. Susanti, B. Mukhopadhyay, An intertwined evolutionary history of methanogenic archaea and sulfate reduction. PLOS ONE 7, e45313 (2012).

35. Y. Liu, M. Sieprawska-Lupa, W. B. Whitman, R. H. White, Cysteine is not the sulfur source for iron-sulfur cluster and methionine biosynthesis in the methanogenic archaeon *Methanococcus maripaludis*. J Biol Chem. 285, 31923–31929 (2010).

36. E. F. Johnson, B. Mukhopadhyay, Coenzyme F_420_-dependent sulfite reductase-enabled sulfite detoxification and use of sulfite as a sole sulfur source by *Methanococcus maripaludis*. Appl Environ Microbiol 74, 3591–3595 (2008).

37. M. Jespersen, A. J. Pierik, T. Wagner, Structures of the sulfite detoxifying F_420_-dependent enzyme from Methanococcales. Nat Chem Biol, 1–8 (2023).

38. J. Zopfi, T. G. Ferdelman, H. Fossing, Distribution and fate of sulfur intermediates— sulfite, tetrathionate, thiosulfate, and elemental sulfur—in marine sediments. In Sulfur Biogeochemistry - Past and Present. v.379 (2004). 10.1130/0-8137-2379-5.97

39. C. Sattler, S. Wolf, J. Fersch, S. Goetz, M. Rother, Random mutagenesis identifies factors involved in formate-dependent growth of the methanogenic archaeon *Methanococcus maripaludis*. Mol Genet Genomics 288, 413–424 (2013).

40. D. G. Gibson, et al., Enzymatic assembly of DNA molecules up to several hundred kilobases. Nat Methods 6, 343–345 (2009).

41. V. Quitzke, J. Fersch, D. Seyhan, M. Rother, Selenium-dependent gene expression in *Methanococcus maripaludis*: Involvement of the transcriptional regulator HrsM. Biochimica et Biophysica Acta (BBA) - General Subjects 1862, 2441–2450 (2018).

42. D. L. Tumbula, R. A. Makula, W. B. Whitman, Transformation of *Methanococcus maripaludis* and identification of a Pst I-like restriction system. FEMS Microbiol Letters 121, 309–314 (1994).

43. B. E. Rubin, et al., Species- and site-specific genome editing in complex bacterial communities. Nat Microbiol 7, 34–47 (2022).

44. L. A. Day, E. L. Kelsey, D. R. Fonseca, K. C. Costa, Interspecies formate exchange drives syntrophic growth of *Syntrophotalea carbinolica* and *Methanococcus maripaludis*. Appl. Environ. Microbiol. 88, e01159–22 (2022).

45. C. Hildenbrand, T. Stock, C. Lange, M. Rother, J. Soppa, Genome copy numbers and gene conversion in methanogenic archaea. J Bacteriol 193, 734–743 (2011).

