## Supplemental materials for "High-throughput genetics enables identification of nutrient utilization and accessory energy metabolism genes in a model methanogen"

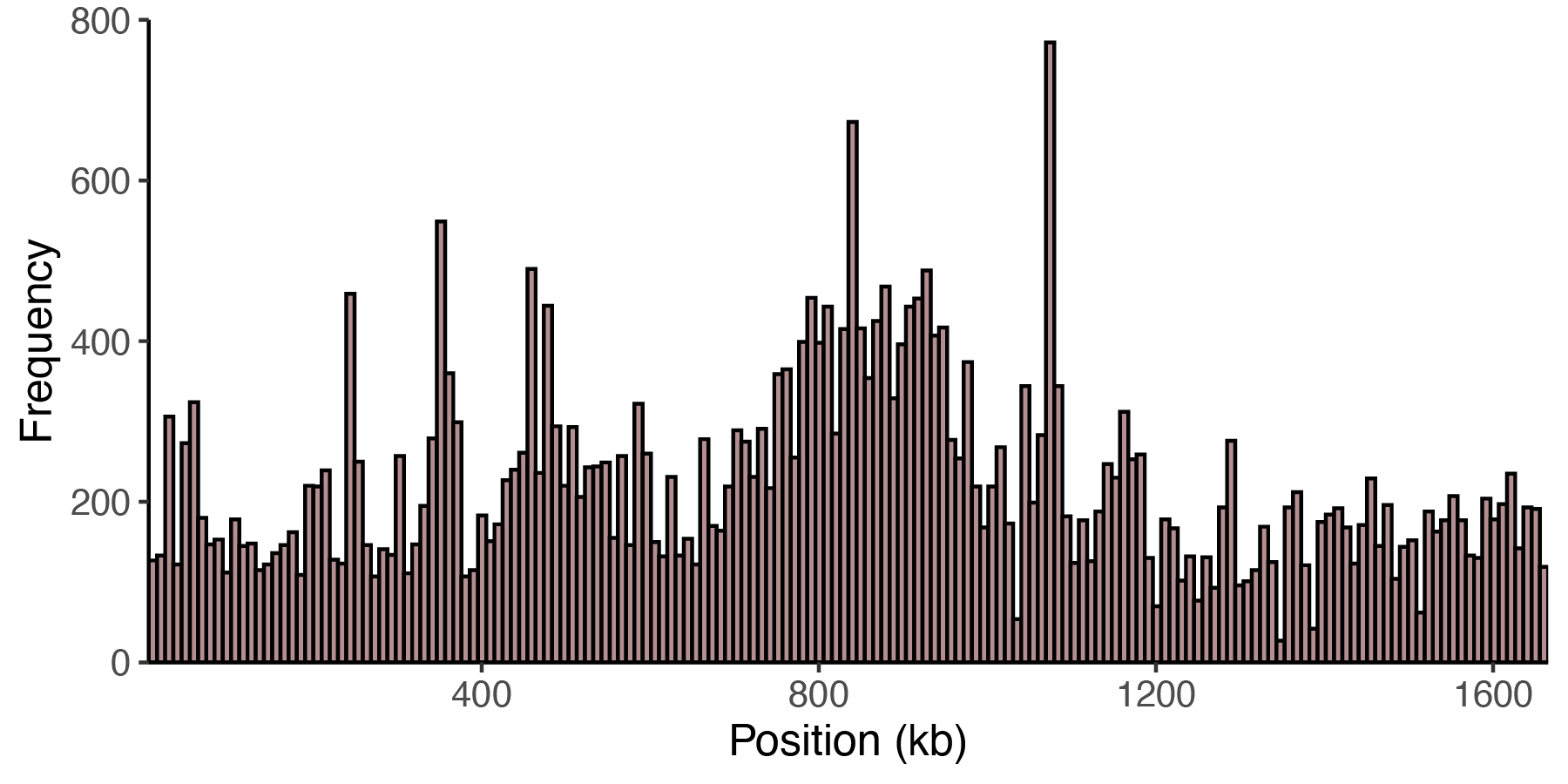
Supplemental Figure 1. Coverage of mapped insertions in the *M. maripaludis* strain S2 RB-TnSeq library across the chromosome.

**Control experiments to account for changes in media from growth medium used to generate the library.**

To minimize the differences between the hydrogen and formate oxidizing medium and increase the buffering capacity for downstream experiments with added compounds we added MOPs buffer to the base H2 oxidizing medium. We also transitioned to using trace minerals and general salts solution with chloride rather than SO_4_ to increase comparability to future experiments not included in this manuscript. We performed fitness assays with the MOPs buffer addition and the SO_4_ free trace minerals There were no specific phenotypes for these medium conditions. Thus, all future experiments were conducted with the addition of MOPS in both the hydrogen and formate oxidizing conditions and SO4 free trace minerals.

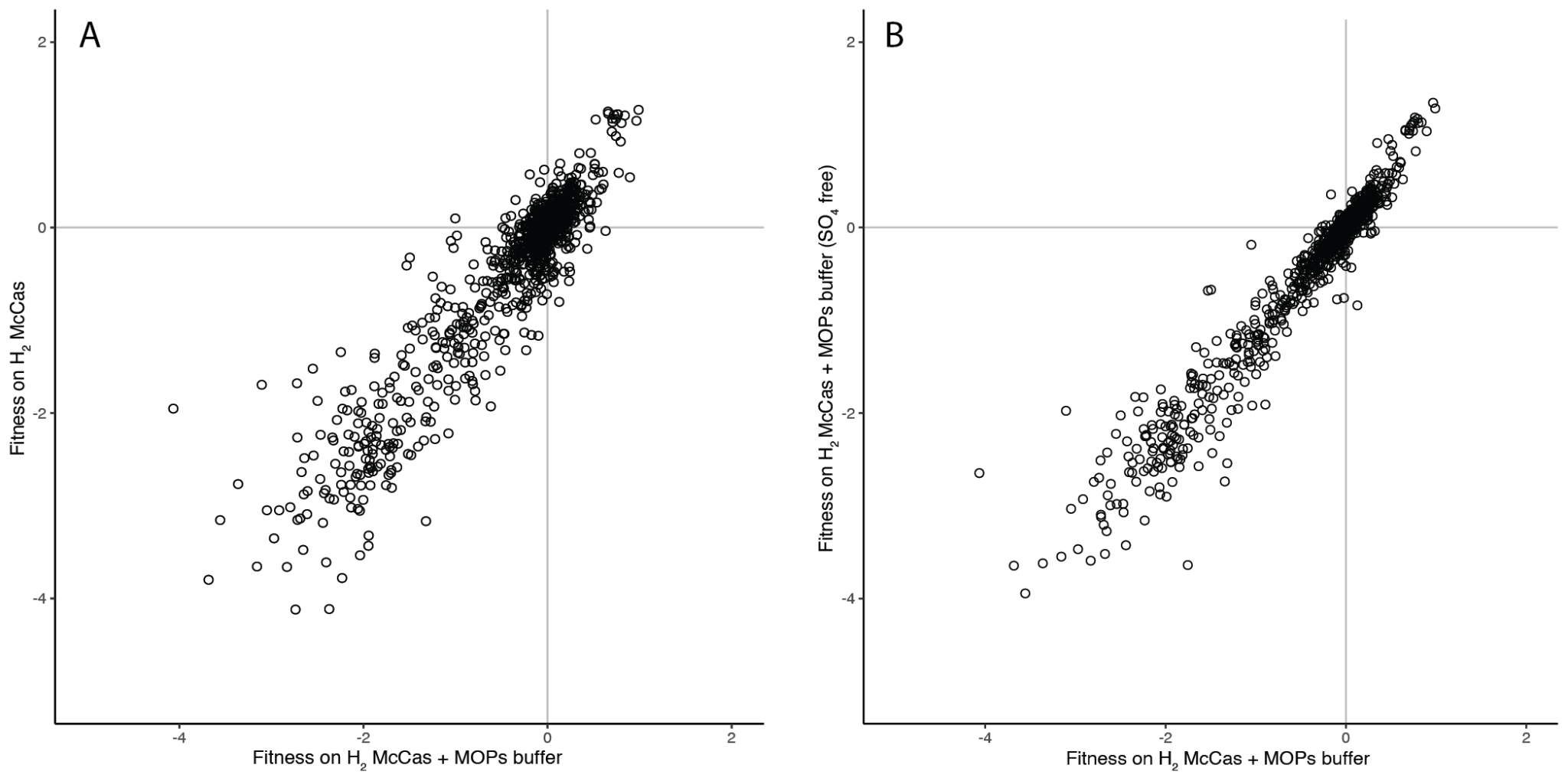

Supplemental Figure 2. Comparison of *M. maripaludis* strain S2 gene fitness in rich medium with addition of (A) MOPs buffer and (B) removal of SO_4_ from trace minerals and general salts solution. Points represent averages of 3 replicates for each condition.

***M. maripaludis* fitness assays conducted in high-throughput in both hydrogen and formate oxidizing conditions**

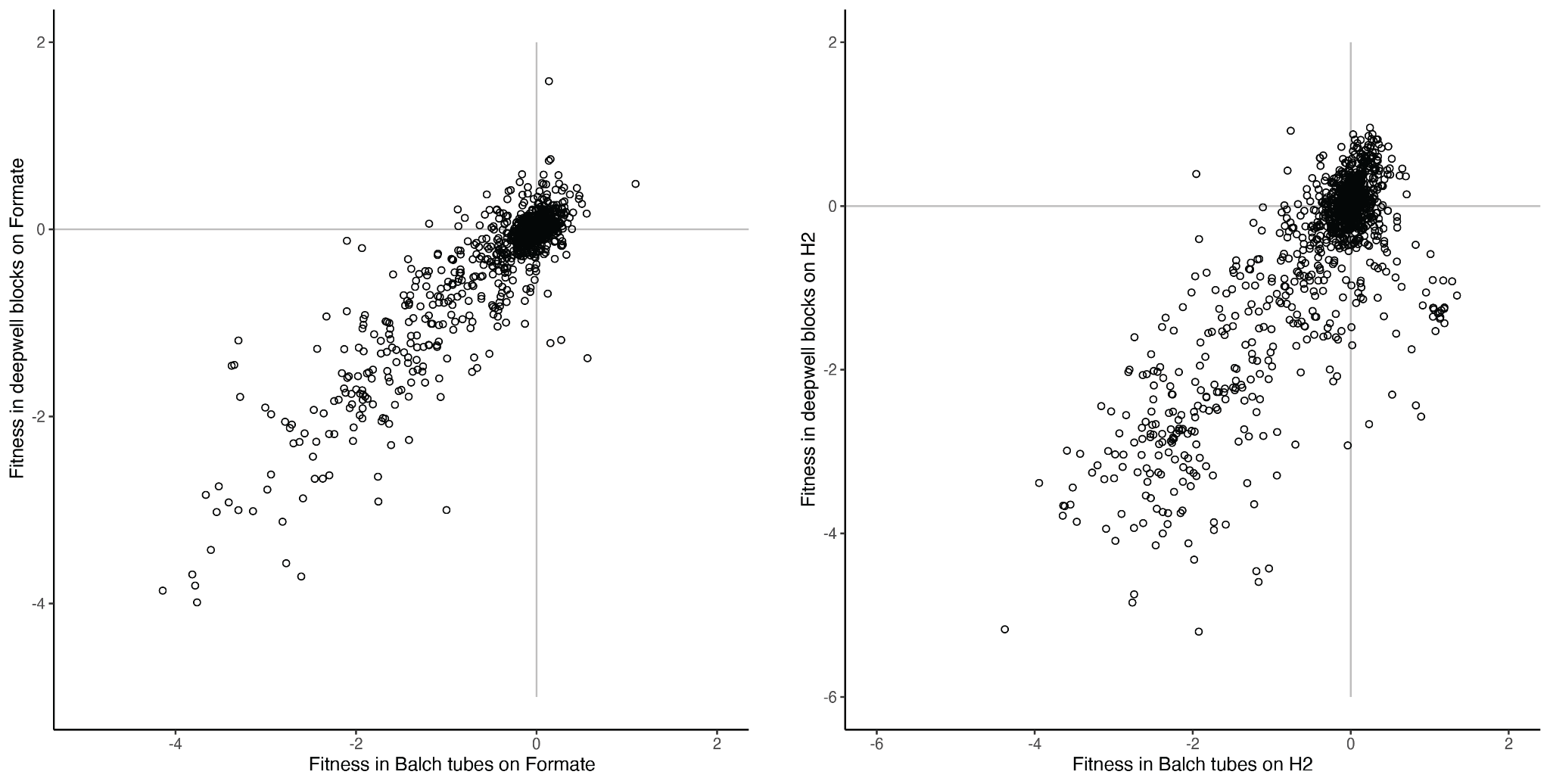

Supplemental Figure 3. Comparison of *M. maripaludis* strain S2 gene fitness data between cultivation techniques using Balch tubes or 96 well deep-well blocks in (A) formate or (B) H_2_ oxidizing conditions. Much of the metabolic characterization done for *M. maripaludis* has been conducted in H_2_ oxidizing conditions. However, to our knowledge the high-throughput characterizations done on hydrogenotrophic methanogens have been in formate oxidizing conditions to circumvent the need to provide high concentrations of H_2_ as an electron donor in a 96 well format (1). We sought to implement a high-throughput method of growth for both hydrogen and formate oxidizing conditions and determine any differences by performing fitness assays with comparable medium in Balch tubes and deep-well blocks. While there are differences in gene fitness between growth with H_2_ in Balch tubes versus deep-well blocks, none were genes of interest in our subsequent experiments. So, we suspect that much of the differences for the H2 oxidizing conditions are a result of differences in H_2_ concentration. Cultures usually reached visible turbidity after 2 days in deep-well blocks compared to 1 in Balch tubes.

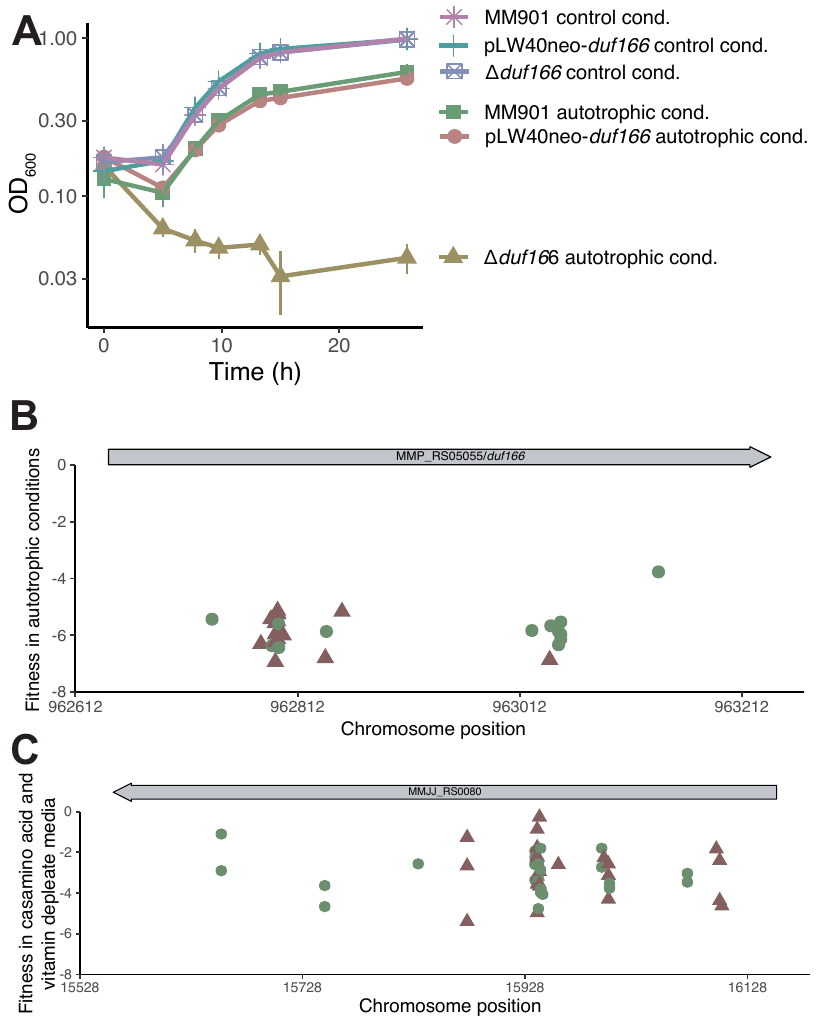

Supplemental Figure 4. Fitness defects in *duf166* (MMP_RS05055). (A) Growth of *M. maripaludis* strain S2 measured in OD_600_ over time for the WT (MM901), deletion strain (*Δduf166*), and complement strain (pLW40neo-*duf166*). Control conditions contain 10 mM acetate and autotrophic conditions lack both casamino acids and acetate. (B) Fitness of individual strains with insertions in the central 10-90% of MMP_RS05055. Fitness values are averages of 3 replicate cultures under autotrophic H_2_ oxidizing conditions (C) Fitness of insertions in the central 10-90% of MMJJ_RS00080. Insertions with the puromycin resistance marker's promoter on the + strand are indicated by green circles. Fitness values are averages of 3 replicate cultures grown in media lacking casamino acids and vitamins under H_2_ oxidizing conditions.

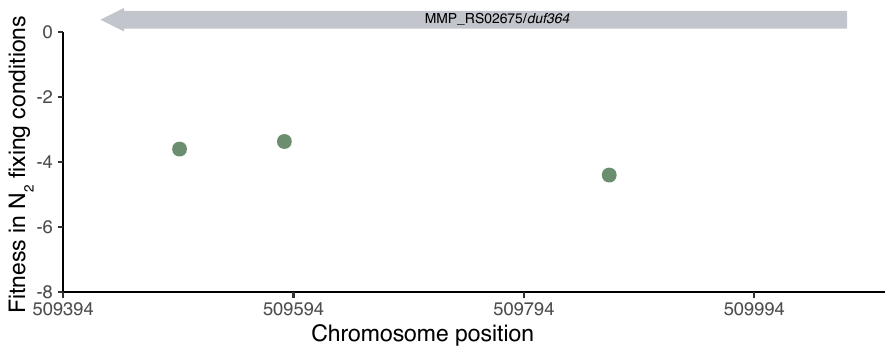

Supplemental Figure 5. Insertions in the central 10-90% of gene MMP_RS02675 which encodes DUF364. Insertions with the puromycin resistance marker's promoter on the + strand are indicated by green circles. Fitness values are averages of triplicate N_2_ fixing & H_2_ oxidizing fitness assays.

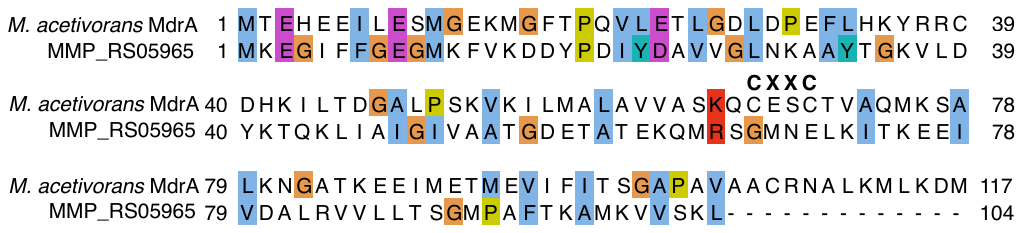

Supplemental Figure 6. Alignment of putative oxygen resistance protein (MMP_RS05965) and *Methanosarcinia acetivorans* MdrA (MA_3736). Conserved cysteine residues (CXXC) essential for MdrA’s disulfide reductase activity (2) are indicated. Amino acids are colored by Clustal X color scheme.

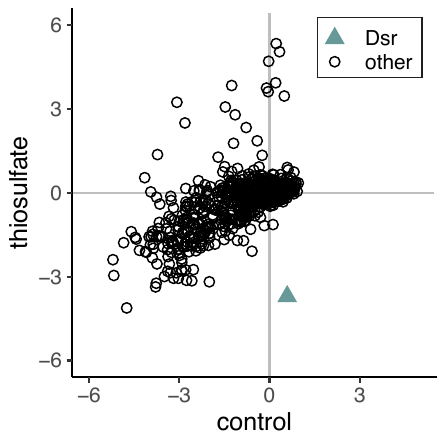

Supplemental Figure 7. Comparison of fitness data between control and 10 mM thiosulfate stress. Points are averages of 3 replicate cultures. Insertions in MMP_RS00450 *dsr* are indicated by a blue triangle and all other genes are represented by open circles.

Supplemental Table 1. Strains and Constructs

| **Strain** | **Description** | **Source** |
| --- | --- | --- |
| *M. maripaludis* strain S2 | Wild-type *Methanococcus maripaludis* strain S2 | (4) |
| *M. maripaludis* strain JJ | Wild-type *M. maripaludis* strain JJ | William Whitman |
| MM901 | *M. maripaludis* S2 Δ*upt* (MMP_RS03565) | (5) |
| KC116/  pLW40neo | MM901 with an empty pLW40neo derivative | (3) |
| KC151/  pLW40neo-*dsr* | Dsr (MMP_RS00450) expression strain | This study |
| KC180/Δ*duf166* | DUF166 MMP_RS05055 deletion strain | This study |
| KC183/  pLW40neo-*duf166* | DUF166 (MMP_RS05055) expression strain | This study |
| **Plasmid** |  |  |
| pJJ605 | Non-replicating plasmid (in *M. maripaludis*) containing mini-mariner transposon and puromycin selection cassette | (6) |
| pT7tnp | *HimarI* transposase expression plasmid | (6) |
| pLD026 | Barcoded version of pJJ605 | This study |
| pLW40neo | Replicative plasmid containing kanamycin/neomycin and ampicillin resistance cassettes | (7) |
| pLW40neo-*dsr* | Replicative plasmid pLW40neo with MMP_RS00450 under control of *Methanococcus voltae* histone promoter | This study |
| pCRuptNeo-Δ*duf166* | Plasmid for the deletion of DUF166 encoded by MMP_RS05055 | This study |
| pLW40neo-*duf166* | Replicative plasmid for complementation of DUF166 MMP_RS05055 under control of *Methanococcus voltae* histone promoter into strain KC180 | This study |

Supplemental Table 2. Primers

| **Oligo name** | **Sequence (5’-3’)** | **Notes** |
| --- | --- | --- |
| 107-pLD26_F | CAGGAGCGGATCGGGAATTAATTgctgctggtgaaagagacc | Gibson assembly primer homology to pJJ605 at BstBI cut site and 109-pLD26_insert1 |
| 108-pLD26_R | GCATGCTCGAGCGGCCcagtgtgatggatataacag | Gibson assembly primer homology to insert2 and pJJ605 at NotI cut site |
| 109-pLD26_insert1 | gctgctggtgaaagagaccctatcttacctgctaaaatctaagttaattactaatttattattaatttattattagattgggcaaaatagtaaaagaaaactaaaggaaacctaatatggtttcctttttttatatatttttaattcactgggggggaatcatttgaaggttggtactatataaaaataatatg | Restore region between cut sites so only adding the U1 20N U2 not also removing 220bp |
| 110-pLD26_insert2 | ctttttttatatatttttaattcactgggggggaatcatttgaaggttggtactatataaaaataatatgcatttaatactagcgacgccatctat*GATGTCCACGAGGTCTCT*NNNNNNNNNNNNNNNNNNNN*CGTACGCTGCAGGTCGAC*gtgtcagaccggggacttatcagccaacctgttatatccatcacactg | Italics represent the U1 and U2 priming sites used for PCR amplification of the barcodes (20 N’s). Underlined is the inverted repeat (IR) |
| DF1-351-Δ*duf166*-upF | GgcgaattgggccctctagaTgagtaacagatcttggtaaaaata | Amplification of 500bp upstream of MMP_RS05055 for markerless deletion of *duf166* in *M. maripaludis* strain S2 |
| DF1-352-Δ*duf166*-upR | Ttattcaccagatactttcattttttcacaagtttagtttaaatca |  |
| DF1-353-Δ*duf166*-dnF | Atgaaagtatctggtgaataaatgaagattgcaattactggaaaag | Amplification of 500bp downstream of MMP_RS05055 for markerless deletion of *duf166* in *M. maripaludis* strain S2 |
| DF1-354-Δ*duf166*-dnR | atcacactggcggccgccttgcttccgaaacaataatcatta |  |
| 156-DUF166F | aactaataggtgaaatgcatgAAAGTAGTTGTAGTTTCTGACGG | Amplification of *duf166* MMP_RS05055 for Gibson assembly into pLW0neo |
| 157-DUF166R | ctacagatctcctaggcgcgccTTATTCACCAGATTTCTTTTTTAATGC |  |
| 142-DsrF | aactaataggtgaaatgcatgaacaaaaatgaaatagccaatctg | Amplification of *dsr* MMP_RS00450 for Gibson assembly into pLW40neo |
| 143-DsrR | acagatctcctaggcgcgccttaataatcttttaatttttcttttaaagaac |  |

**REFERENCES**

1.	Weimar MR, Cheung J, Dey D, McSweeney C, Morrison M, Kobayashi Y, Whitman WB, Carbone V, Schofield LR, Ronimus RS, Cook GM. 2017. Development of Multiwell-Plate Methods Using Pure Cultures of Methanogens To Identify New Inhibitors for Suppressing Ruminant Methane Emissions. Applied and Environmental Microbiology 83:e00396-17.

2.	Lessner DJ, Ferry JG. 2007. The Archaeon Methanosarcina acetivorans Contains a Protein Disulfide Reductase with an Iron-Sulfur Cluster. J Bacteriol 189:7475–7484.

3.	Day Leslie A., Kelsey Elisa L., Fonseca Dallas R., Costa Kyle C. 2022. Interspecies Formate Exchange Drives Syntrophic Growth of Syntrophotalea carbinolica and Methanococcus maripaludis. Applied and Environmental Microbiology 88:e01159-22.

4.	Whitman WB, Shieh J, Sohn S, Caras DS, Premachandran U. 1986. Isolation and characterization of 22 mesophilic methanococci. Syst Appl Microbiol 7:235–240.

5.	Costa KC, Wong PM, Wang T, Lie TJ, Dodsworth JA, Swanson I, Burn JA, Hackett M, Leigh JA. 2010. Protein complexing in a methanogen suggests electron bifurcation and electron delivery from formate to heterodisulfide reductase. PNAS 107:11050–11055.

6.	Sattler C, Wolf S, Fersch J, Goetz S, Rother M. 2013. Random mutagenesis identifies factors involved in formate-dependent growth of the methanogenic archaeon Methanococcus maripaludis. Mol Genet Genomics 288:413–424.

7.	Dodsworth JA, Leigh JA. 2006. Regulation of nitrogenase by 2-oxoglutarate-reversible, direct binding of a PII-like nitrogen sensor protein to dinitrogenase. PNAS 103:9779–9784.
